# Refrigerated storage and cryopreservation of hormone induced sperm in a threatened frog

**DOI:** 10.1101/2023.07.15.548973

**Authors:** Rose Upton, Natalie E. Calatayud, Simon Clulow, Darcie Brett, Alana L. Burton, Kim Colyvas, Michael Mahony, John Clulow

## Abstract

There are strong potential benefits of incorporating assisted reproductive technologies (ARTs) into conservation programs for the management of threatened amphibians as the global amphibian decline continues. As sperm cryopreservation and other ARTs advance in common species, focus on non-lethal sperm collection methods for threatened amphibians is imperative. We aimed to realise this goal by testing various doses of exogenous hormones for non-lethal induction of spermiation in a threatened frog (*Litoria aurea*) and develop cold storage and cryopreservation protocols following the recovery of urinic sperm. Our major findings include: (1) that sperm release could be induced in high concentrations with 20 IU/g bodyweight of human chorionic gonadotrophin (hCG); (2) high levels (>50%) of live, motile sperm could be recovered post-cryopreservation by treating the sperm with 15% v/v DMSO and 1% w/v sucrose pre-freeze; and (3) urinic sperm stored at 5°C retained motility over a 14-day period. Our findings demonstrate that it is possible to obtain and store large quantities of quality sperm from a threatened amphibian via non-lethal means, representing an important step forward for the use of ARTs in conservation programs for rare and threatened species.

**Lay Summary:** Amphibians are undergoing an extinction crisis unparalleled in any other taxa. The development of assisted reproductive technologies, such as sperm cryopreservation and IVF have an important role to play in the conservation management of amphibians globally. Here we report non-lethal methods of sperm collection and storage in a highly threatened amphibian. Major findings include successfully induced sperm release in high concentrations, retention of ∼50% live, motile sperm after 7 days storing it at 5°C, and successful recovery of of more than 50% live, motile sperm following cryopreservation. Our results demonstrate the viability of obtaining and storing sperm from rare and threatened species via non-lethal means, providing an important step forward for the use of ARTs in conservation programs around the globe.

## 1. Introduction

Amphibians are subject to threatening processes that are reducing population sizes below critical thresholds for the retention of genetic diversity, to levels that threaten their persistence through inbreeding, and loss of heterozygosity and allelic richness. One example of growing significance is climate change. This increasingly threatens to drive amphibian extinction events by altering climate driven habitat alteration and an increased frequency and intensity of fire in the landscape [1-3]. As species decline, there is a pressing need for strategies that protect the genetic diversity of small and isolated populations. Assisted reproductive technologies (ARTs), including gamete induction through exogenous hormones, sperm cryopreservation and extended cold storage of sperm without cryopreservation, have the potential to bolster efforts to manage the most serious aspects of genetic loss in declining populations [4-9]. Sperm cryopreservation and biobanking can be utilised to manage genetics effectively through storage of founder genomes and the reduction of costs of captive breeding programs [6-8]. Nevertheless, especially for threatened species, there is room for improving sperm collection methods so that they are minimally invasive.

Historically, many studies reporting cryopreservation of sperm in amphibians have utilised testicular macerates, involving euthanasia, as a source of good quality, concentrated sperm that can be diluted out to large working volumes [10-13]. While testicular macerates have a place in laboratory studies and conservation programs in the case of opportunistic collection from recently deceased animals, development of non-lethal methods such as hormonal induction of urinic sperm (release of mature sperm cells in urine) is critical when working with threatened species, because they do not harm the adult male and can be repeated several times. Furthermore, sperm acquired non-lethally and cryopreserved (biobanked) can be used to retain heterozygosity in both captive and wild populations [7, 8].

Most reports of hormonal induction of sperm release to date indicate a preference for the use of human chorionic gonadotropin (hCG) for producing concentrated high quality sperm samples in urine (i.e. from Pelodryadidae, Bufonidae and Limnodynastidae families; [14-19], with a smaller number of studies indicating a preference for gonadotropin-releasing hormone (GnRH-a) [16, 20, 21]. While targeting different levels of the hypothalamic-pituitary gonadal axis, both hormones ultimately lead to the release of urinic sperm (for details see [14, 22]). While many studies report some success with exogenous hormones for inducing sperm release in a range of species, subsequent cryopreservation of urinic sperm has only recently been explored as an alternative to testicular macerates [23-28].

An alternative to sperm cryopreservation, which can be difficult in some species, is cold storage. Amphibian sperm, particularly from aquatic breeding temperate species, has a high tolerance to cooling and thermal shock with reports of sperm held at temperatures between 0-5°C for periods of up to 21 days with varying degrees of retained motility and vitality [11, 27, 29-37]. Storing sperm at cool temperatures (0-5°C) can be used as a standalone procedure, or as an intermediary step towards cryopreservation as an endpoint [31]. Cold storage of sperm has the potential to allow collection, and transport of sperm from wild populations in instances where the application of cryopreservation technologies *in situ* is logistically hard to implement.

We aimed to develop a suite of ART protocols for a threatened frog, with application to conservation programs in mind. This included (1) to test the efficacy of hCG (20 and 40 IU/g bodyweight) and GnRH-a (0.25 and 0.5 μg/g bodyweight) in the induction of urinic sperm; (2) to test previously developed cryopreservation protocols to optimise post-thaw recovery of hormonally induced urinic sperm at 0- and 30-minutes post-thaw; and (3) to test the rate of decay in motility of urinic sperm held in cold storage (5°C), without cryopreservation, over a 14-day period.

## 2. Methods

### 2.1 Study Species

We investigated our study aims using a model threatened frog, *Litoria aurea. L. aurea* is an Australian frog that has undergone range contractions of more than 90% since the 1980s and is now considered nationally vulnerable [38]. Due to the ease of which the species is able to be kept in captivity, it has become a model for developing ARTs in threatened amphibians in the past, including the recent production of reproductively viable young through IVF using opportunistically collected cryopreserved testicular sperm [39].

### 2.2 Husbandry

Male *L. aurea* from the University of Newcastle breeding program were housed in plastic terraria (30 x 18 x 20 cm), with 25% terrestrial environment (autoclaved pebbles) and 75% aquatic environment (aged tap water), for the duration of the study (May and June 2020). Plastic aquaria plants were provided as refuges. Live crickets supplemented with calcium and vitamin powder (Multical Dust; Vetafarm, Wagga Wagga, NSW, Australia) were provided as food twice weekly for the duration of the experiment.

All animals underwent a precautionary protocol of heat treatment to ensure no chytridiomycosis infection was present. Animals were placed in UV- and temperature-controlled cabinets (TRISL – 1175, Thermoline Scientific Equipment, Wetherill Park, NSW) on a 12:12 UV light cycle started at 25°C. Temperature was increased at 2°C per day until 37°C was reached, where the temperature was held for 6 hours. This temperature has been shown to kill *Batrachochytrium dendrobatidis* within 4 hours (Johnson et al., 2003). Subsequently, temperature was reduced at 2°C per day until the original 25°C was reached. Animals were kept at 25°C until the conclusion of the experiment in June 2020. While experiments were conducted across the shoulders of the *L. aurea* breeding season (May - June), all males included in experiments were in season, confirmed by the presence of darkened nuptial pads.

All experiments were carried out in accordance with International, National and Institutional standards for the care and welfare of animals and in accordance with the University of Newcastle’s ethics approval, A-2013-328. Animals were collected and held under NSW Scientific Licence SL101269.

### 2.3 Experiment 1: Effect of hCG and GnRH-a doses on spermiation response of L. aurea

In order to determine the effect of hCG (Chorulon®, Intervet) and GnRH-a (des-Gly^10^, D-Ala^6^, Pro-LHRH, Bachem) dose on the spermiation response of *L. aurea*, 20 males (n=5 per treatment) were assigned to one of 4 experimental treatments: (1) 20 IU/g body weight hCG; (2) 40 IU/g of body weight hCG; (3) 0.25 μg/g body weight GnRH-a, and; (4) 0.5 μg/g body weight GnRH-a, across 10 days (2 experimental treatments per day) between 6th May – 18th June 2020. As animals were originally sourced from wild populations, age was unknown (however males reach sexual maturity at around 1 year of age; the weight range of males used in the study was 14.3 ± 2.3 g (mean and standard error, n=20).

Prior to injection, a urine sample was collected from each male and found to be aspermic in all males used in the experiment. Males were weighed to the nearest 0.1 g for the determination of hormone dose to be administered by body weight. Hormones were diluted in simplified amphibian ringer (SAR; 113 mM NaCl, 1 mM CaCl_2_, 2 mM KCl, 3.6 mM NaHCO_3_; ∼220 mOsm/kg; recipe [10]) to a final volume of 200 μl. In addition, each day animals were treated, a male (total, n=10) was administered with a control treatment of 200 μl of SAR to test for spermiation in negative controls. Hormones were administered subcutaneously via the dorsal lymph sac using a 31-gauge insulin syringe (BD, New Jersey, United States).

Once treated, frogs were placed in individual plastic containers (15 x 7 x 10 cm) with ∼1 cm of water in the bottom to ensure hydration. At allotted collection intervals (1, 2, 3, 4, 5, 6, 24, 48 hours post-injection), a thin plastic gel-loading tip (Corning, New York, United States), was used as a catheter to collect a spermic urine sample. The pipette was gently placed ∼3-5 mm into the cloaca and gently moved in and out to facilitate urine collection by capillary action. The sample was transferred to a 0.5 mL Eppendorf tube and was kept on ice until volume, motility and sperm concentration could be determined (within the hour).

Concentration (cells/mL) of sperm in urine samples were assessed with an Improved Neubauer haemocytometer counting chamber at each collection interval. Ten microlitres was pipetted into each of two chambers, and the number of sperm in at least 5 secondary quadrats (per chamber) was counted and used to calculate total sperm per millilitre. For concentrated samples, sperm were diluted, as necessary, with SAR and the dilution factor was taken into account in the final calculation. For particularly dilute samples, all primary quadrats of the counting chamber were counted. The number of either primary or secondary quadrats was taken into account for the final calculation. The total number of sperm per sample was calculated using the volume of sample collected.

Due to small volumes of urinic sperm available for assessment, motility in samples was analysed after a 1:5 dilution in SAR. While this was found to be less favourable than analysis of undiluted samples, it allowed sufficient volume for assessment of the effect of hormone type and dosage rate on motility where only small volumes were available. Sperm were categorised as either non-motile, or motile, and at least 100 sperm were counted at 400x magnification across at least four fields of view using a Kyowa Unilux-12 microscope and phase contrast optics. Due to small sample volumes, only one replicate count could be performed per sample.

### 2.4 Experiment 2: Cryopreservation of urinic sperm of L. aurea

A further four males were used to test the efficacy of cryoprotectant solutions previously optimised using sperm derived from testicular macerates (see [39, 40]) on the recovery of urinic sperm of *L. aurea*. Previous studies showed 15% v/v DMSO with 10% w/v sucrose provided sufficient recovery of sperm for fertilisation of ova [39], however a more recent study on related pelodryadid species shows 15% v/v DMSO in combination with 1% w/v sucrose could be a more appropriate cryoprotectant for Australian tree frogs [40]. Thus, both of these cryoprotectants were chosen to test the recovery of urinic sperm.

Males were injected with 20 IU/g body weight hCG and samples collected over 6 hours until an accumulative volume of at least 50 μl was achieved. Forty microlitres of urinic sperm were diluted 1:5 (to a final volume of 240 μl) with the appropriate ice-cold cryoprotectant and loaded into 0.25 μl Cassou straws (Minitube, Smythesdale, Victoria, Australia) at a volume of 120 μl per straw. Two straw replicates were produced per treatment, per animal. Straws were frozen in a programmable freezer (Freeze Control® CL-3300; CryoLogic Pty Ltd, Blackburn, Victoria, Australia), as per [13].

Straws were thawed on a bench at room temperature (∼21°C) and assessed as per [40]. Briefly, motility was assessed by diluting samples 1:9 so that the final concentration of DMSO and sucrose in each sample was equal to 15% v/v DMSO and 1% w/v sucrose. This allowed assessment of motility at equal osmolalities across treatments whilst accounting for any dilution effects on motility. Following dilution, sperm were assessed as per Experiment 1. Sperm vitality (scored as the proportion of cells with intact cell plasma membranes) was assessed with an Eosin-Y dye exclusion assay [39, 40]. Sample was mixed 1:1 with Eosin-Y to a final volume of 10 μl and viewed under a coverslip at 400x magnification. Sperm with clear cytoplasm (excluding the dye) were scored as live and sperm stained pink were scored as dead (membrane permeable to the dye). For both motility and vitality, assessment of each straw counted at least 100 sperm across four fields of view, in duplicate. All sperm collections for this experiment occurred in June 2020, at which time a pre-freeze assessment was made, and post-thaw assessment occurred September 2020, following three months of cryostorage.

### 2.5 Experiment 3: Cold storage of urinic sperm without cryopreservation

To determine the longevity of motility in urinic sperm samples at 5°C, excess non-cryopreserved samples from Experiments 1 and 2 were kept in refrigerated storage for a period of 14 days. Aliquots (15-25 μl) of spermic urine from seven males were kept in 0.5 mL Eppendorf tubes. Where volume permitted (n=3), three aliquots of spermic urine from the same animal were used and motility analysed on days 1, 3, 5, 6, 8, 9, 10, 11, 12, 13 and 14 or until the sample ran out. Otherwise (n=4), a single aliquot was used and analysed on days 1, 2, 3 and 5. Initial motility was also determined on day 0. Motility of urinic sperm was assessed undiluted, with no added media, by adding a 2 μl drop under a 12 mm diameter coverslip (Menzel-Gläser, Saarbrückener, Germany) coverslip at 200x magnification after allowing 1 minute to reach room temperature. While osmolality couldn’t be measured for each sample due to small volumes, a subset of urine samples gave an average osmolality of 50 mOsm/Kg (n=5). Categorisation of motility was as per Experiments 1 and 2. Due to small volumes available, only a single count was completed, per sample, each day. Sperm were aerated by gently bubbling air into the sample with a pipette for about 5-10 seconds every day of storage, regardless of whether motility was being assessed.

### 2.6 Statistics

Generalised Linear Mixed Models (GLMM) were used to fit Poisson regressions for sperm concentration (cells/mL) and total sperm numbers (cells/sample). The raw counts were used with an offset of the log reciprocal of an adjustment factor (based on dilution factor, volume and the number of squares counted as appropriate) to convert the raw counts to cells/mL or cells/sample respectively. Treatment and hour were utilised as main effects and animal ID as a random effect. Using AIC as a model fit criterion, it was found the model needed to be expanded to account for zero inflation as a function of time (hours) to account for excess zeros beyond those expected from the Poisson distribution and use the negative binomial distribution to account for overdispersion. Significance was determined by pairwise comparisons to determine ratios of expected counts between conditions and 95% confidence intervals.

To determine the effect of the four hormone treatments on motility, we used Generalised Linear Mixed Models (GLMM) to fit binary logistic regressions (interpreted as the proportion of successful cases (i.e. motile sperm) with total number of sperm equalling the weights for the model. Treatment and hour were used as main effects and overdispersion was addressed using an observation level random effect. The interaction between treatment and hour was not significant and was dropped from the model.

To determine the effect of the two cryoprotectants and time post-thaw on motility and vitality, we used GLMMs to fit binary logistic regressions (interpreted as the proportion of successful cases (i.e. motile or live sperm) with total number of sperm equalling the weights for the model. We used a 2x2 factorial model design with cryoprotectant (15% v/v DMSO with either 1 or 10% w/v sucrose) and time post-thaw (0 and 30 minutes) as main effects and overdispersion was addressed using an observation level random effect. The interaction between cryoprotectant and time post-thaw was not significant and was dropped from the model.

To determine the effect of the days in cold storage (without cryopreservation) on motility, we used generalised linear model to fit a binary logistic regression (interpreted as the proportion of successful cases (i.e. motile) with total number of sperm equalling the weights for the model. The number of days in storage was used as the model main effect (continuous) due to overdispersion in the model, the GLM was run in the quasibinomial distribution.

All analyses were completed using R (Version 3.6.2) with R packages glmmTMB used for zero-inflated negative binomials and lme4 used for all other GLMM modelling [41, 42]. The package emmeans was used to model estimated marginal means (EMMs) and 95% confidence limits were calculated and back transformed to sperm concentrations (in cells/mL and cells/sample) and motility and vitality proportions. Ratios (for sperm concentration data) and odds ratios (for proportion data) comparing treatments were also generated using the package emmeans [43]. All graphing was completed using ggplot2 and gridExtra [44, 45].

## 3. Results

### 3.1 Experiment 1: Effect of hCG and GnRH-a doses on spermiation response of L. aurea

There was a significant effect of treatment and collection time on sperm concentration (likelihood ratio test (LRT) χ^2^(3) = 15.0, p < 0.01 and LRT χ^2^(7) = 43.1, p < 0.001 for treatment and collection time respectively) and total sperm numbers per sample (LRT χ^2^(3) = 13.2, p < 0.01 and LRT χ^2^(7) = 77.7, p < 0.001 for treatment and collection time respectively). The superior hormone was hCG across all timepoints with a peak in sperm concentration at 3 hours and total sperm numbers at 2 hours. Ratios and 95% confidence intervals were generated to further compare the differences in treatments, with larger ratios indicating bigger differences in spermiation response.

For sperm concentration, both hCG treatments were significantly better than the two GnRH-a treatments with the ratio of expected sperm concentrations ranging from the lowest for the comparison 40 IU/g hCG and 0.25 μg/g GnRH at 11.2, 95% CIs: [1.8, 70.3] to the highest for 20 IU/g hCG and 0.5 μg/g GnRH at 36.7, 95% CIs: [5.8, 232.3] (see Supplementary Table 1 for ratios and 95% CIs). Likewise, for total sperm numbers, hCG outperformed GnRH-a, with the ratio of expected sperm concentrations ranging from the lowest for the comparison 40 IU/g hCG and 0.25 μg/g GnRH at 13.8, 95% CIs: [2.2. 85.5] to the highest for 20 IU/g hCG and 0.5 μg/g GnRH at 20.3, 95% CI [3.3, 126.0] (Supplementary Table 2).

Predicted sperm concentration did not exceed 1.0x10^7^ cells/mL in 0.25 μg/g GnRH-a or 3.91x10^6^ cells/mL in 0.5 μg/g GnRH-a but reached as high as 1.44x10^8^ and 1.12x10^8^ cells/mL in 20 IU/g and 40 IU/g hCG respectively. Each of these maximum sperm concentrations occurred 3 hours post-injection (Figure 1a) and was significantly higher than sperm collected from 5 hours onwards (see Table S3 for ratios and 95% CIs). Despite high concentrations, total volumes collected were low, with no statistically significant difference between treatments (F(3,17)= 1.74, p = 0.20). Total volumes collected ranged from 136.2 ± 28.8 to 400.5 ± 134.9 μl, with the highest volumes of urine in the treatments with the lowest concentrations (Table 1). Predicted total sperm numbers peaked at 2 hours post-injection for all treatments, with 2.31x10^5^ and 1.78x10^5^ cells/sample for 0.25 μg/g GnRH-a and 0.5 μg/g GnRH-a respectively and 3.61x10^6^ and 3.19x10^6^ cells/sample in 20 IU/g and 40 IU/g hCG respectively (Figure 1b). The total sperm numbers derived at 2 hours was significantly higher than sperm numbers collected from 4 hours onwards (see Supplementary Table 3 for ratios and 95% CIs).

**Figure 1.**
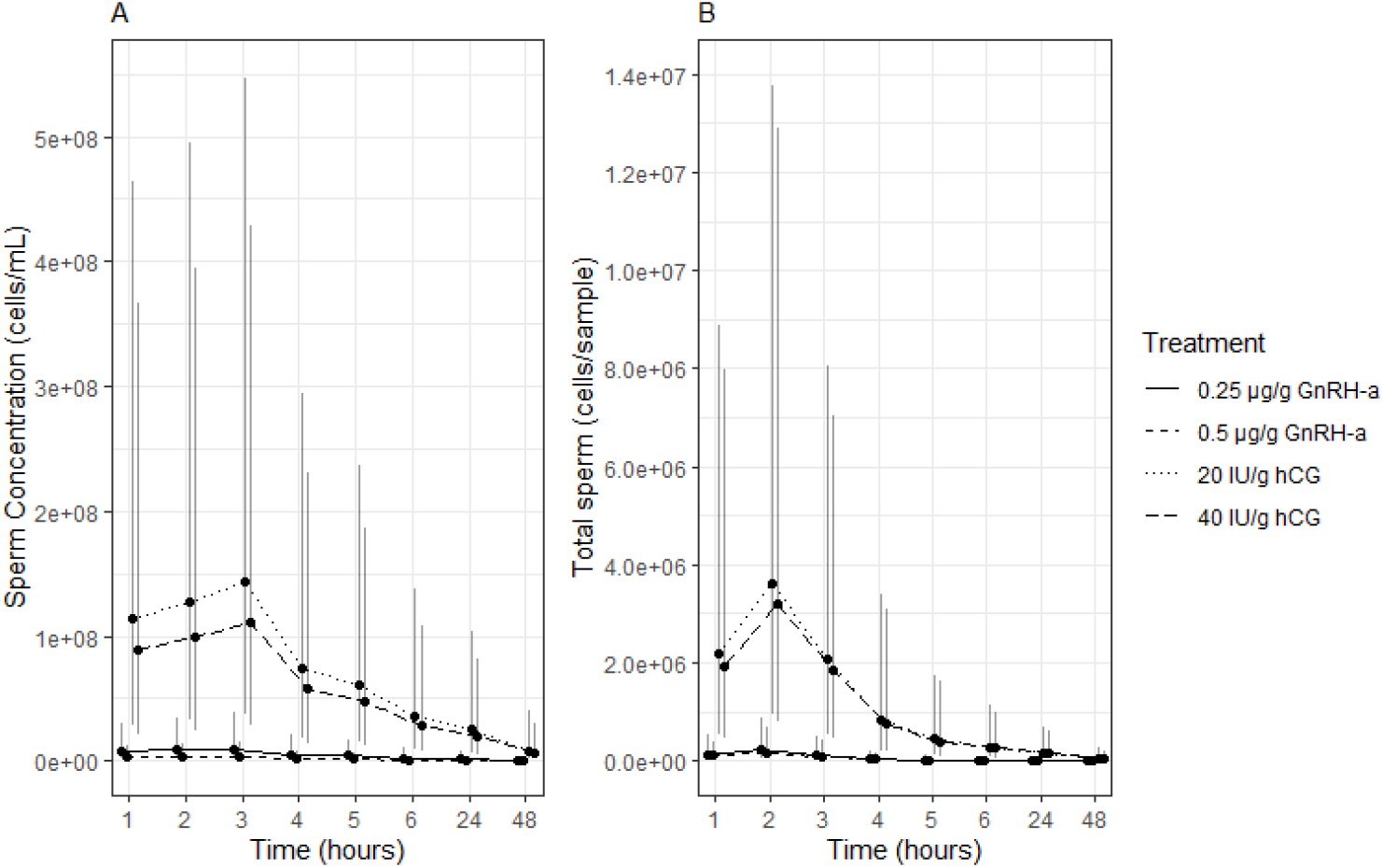
Effect of treatment on induction of spermiation in Litoria aurea (n=5 per treatment) for (a) sperm concentration (cells/mL) and (b) total sperm numbers (cells/sample) over a 48-hour collection period. Zero-inflated negative binomial GLMM; black dots represent estimated marginal means and error bars equal 95% confidence intervals.

**Table 1.** Average volume of urinic sperm (± standard error) collected per hormone treatment in Litoria aurea (n=5 per treatment).

| Treatment | Volume (μl) |
| --- | --- |
| 0.25 μg/g GnRH-a<sup>1</sup> | 219.4 ± 69.1 |
| 0.5 μg/g GnRH-a | 400.5 ± 134.9 |
| 20 IU/g hCG<sup>2</sup> | 193.3 ± 78.0 |
| 40 IU/g hCG | 136.2 ± 28.8 |

<sup>1</sup>Gonadotropin-releasing hormone
<sup>2</sup>Human chorionic gonadotropin

There was a significant effect of treatment (LRT χ^2^(3) = 20.3, p < 0.001), but not time (LRT χ^2^ (5) = 6.7, p = 0.24) on the motility of sperm, with 20 IU/g hCG yielding sperm of significantly higher motility averaged across all time-points than all other treatments (EMM= 60.7%, 95% CIs: [43.6, 75.6]; see Table S4 for ratios and 95% CIs). 40IU/g hCG had a much lower predicted motility (EMM= 23.7%, 95% CIs: [13.3, 38.7]), which was not significantly different to either GnRH-a treatments (see Supplementary Table 4). Of the GnRH-a treatments, 0.25 μg/g had higher motility (EMM= 30.8%, 95% CIs: [17.2, 48.8]) than 0.5 μg/g (EMM=11.7%, 95% CIs: [5.7, 22.6]) which was significantly higher (Figure 2, Supplementary Table 4).

**Figure 2.**
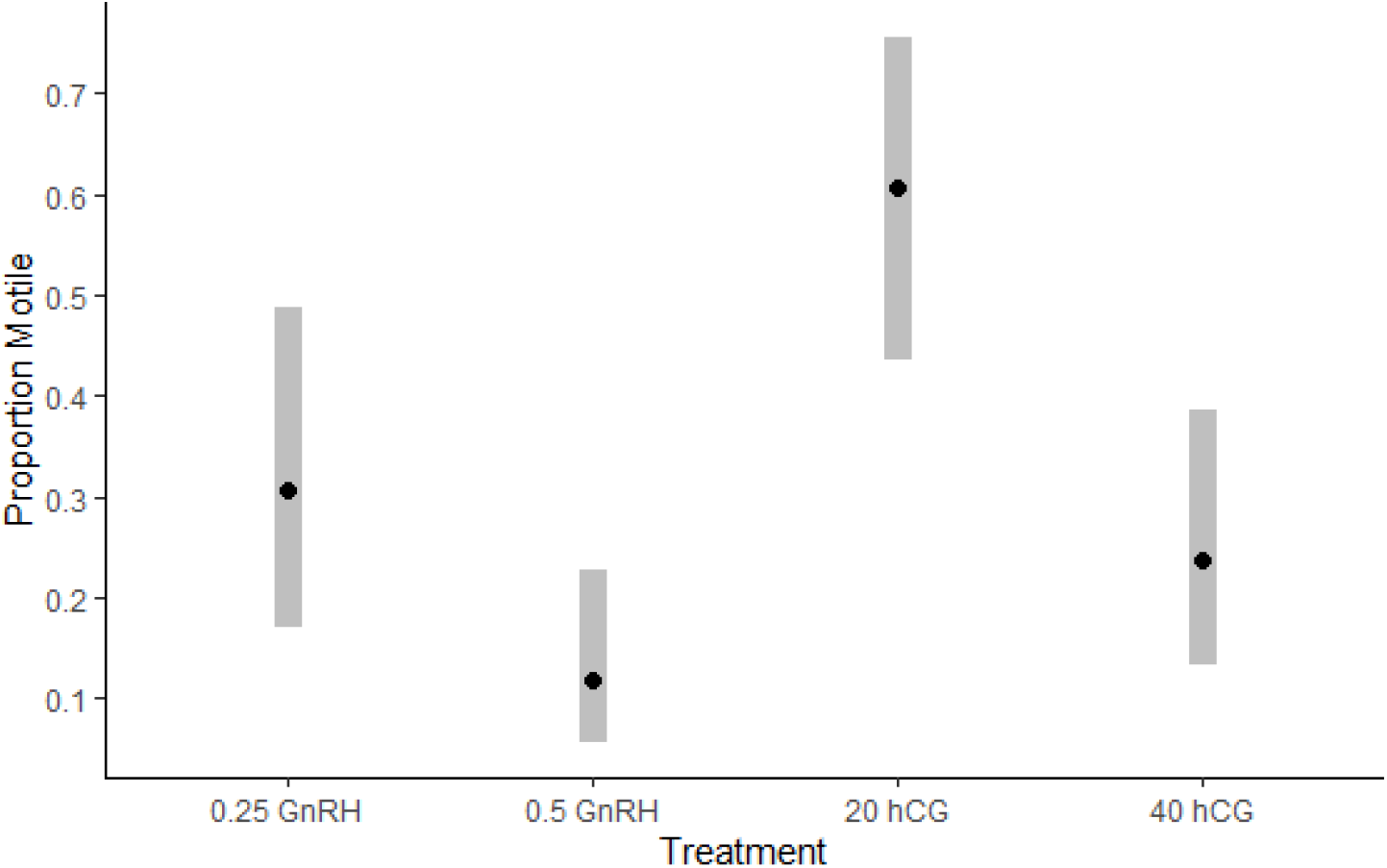
Predicted motility for sperm induced by each treatment (0.25 μg/g body weight GnRH-a, 0.5 μg/g body weight GnRH-a, 20 IU/g bodyweight hCG, 40 IU/g body weight hCG) in Litoria aurea (n=5 per treatment). Estimated marginal means determined by generalised linear mixed modelling. Error bars = 95% confidence intervals.

### 3.2 Experiment 2: Cryopreservation of urinic sperm of L. aurea

There was a significant effect of sucrose concentration on motility (likelihood ratio test (LRT) χ^2^(1) = 31.9, p < 0.001) and vitality (LRT χ^2^(1) = 23.1, p < 0.001), but not time post-thaw (LRT χ^2^(1) = 1.8, p = 0.2 & LRT χ^2^(1) = 0.9, p = 0.3 for motility and vitality respectively). Fresh sperm before addition of cryoprotectants was higher in motility (EMM: 82.0%, 95% CIs: [60.1, 93.2]) and vitality (EMM: 75.3%, 95% CIs: [59.6, 86.3]) than all post-thaw parameters. Odds Ratios (OR) and (95% confidence intervals were used to further compare treatments.

At both 0- and 30-minutes post-thaw, 15% v/v DMSO with 1% w/v sucrose achieved greater recovery of motility and vitality than when 10% w/v sucrose was used instead. For the best cryoprotectant, 15% v/v DMSO with 1% w/v sucrose, there was a slight decrease in motility from 52.7% to 42.4%, and in vitality from 53.1% to 48.8% 30 minutes post-thaw, though this was not significant (Figure 3). Likewise, when using 15% v/v DMSO with 10% sucrose, there was a slight decrease in motility from 14.0% to 9.8% and in vitality from 30.1% to 26.6% 30 minutes post-thaw (Figure 3). Urinic sperm cryopreserved with 15% v/v DMSO with 1% w/v sucrose were almost 7 times more likely to recover motility (OR: 6.9, 95% Confidence intervals (95% CI) [3.8, 12.4]) and 2.6 times more likely to recover vitality (OR: 2.6, 95% CI [1.8, 3.8]) across both time points.

**Figure 3.**
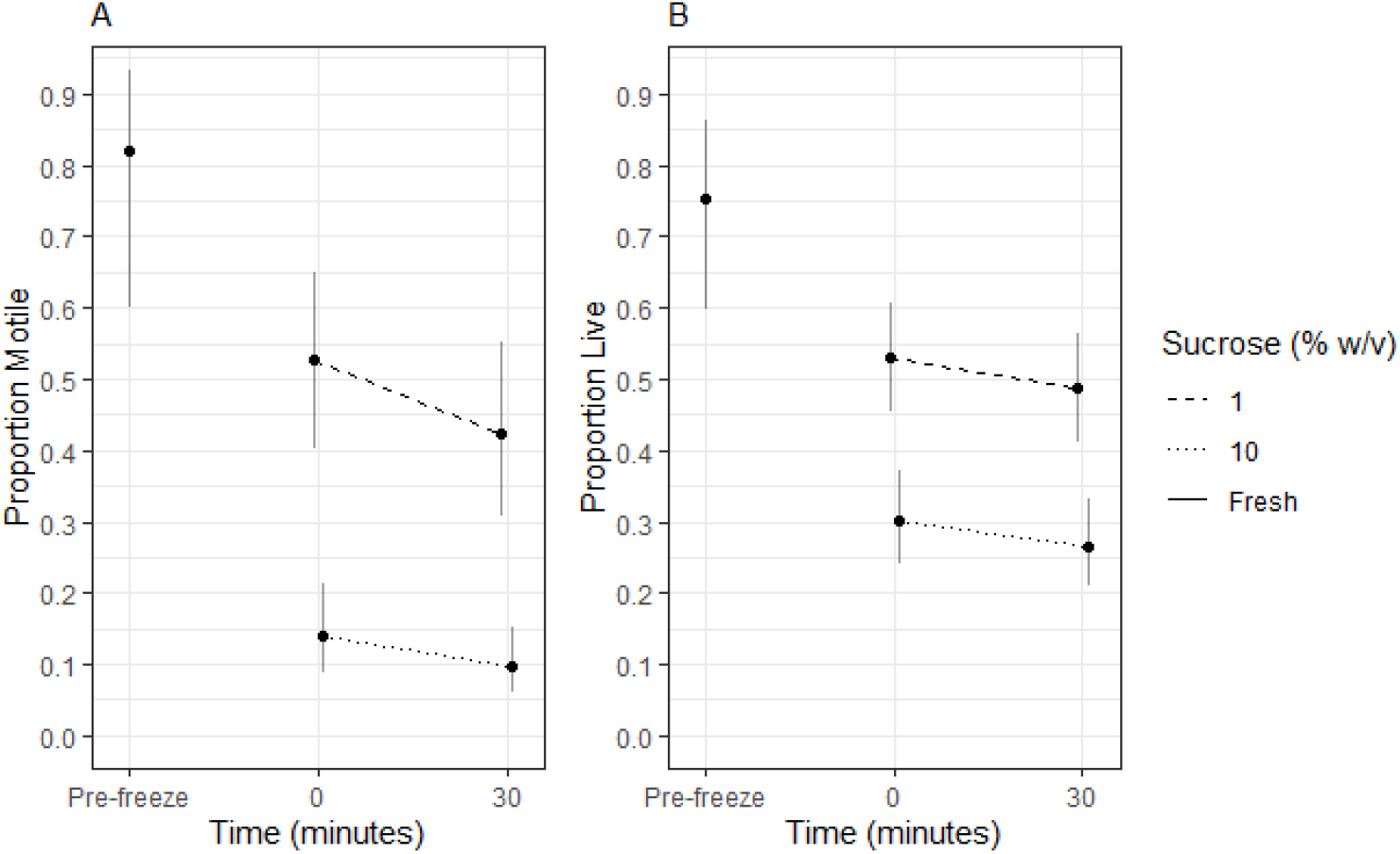
Predicted motility and vitality for pre-freeze (cryoprotectant-free) urinic sperm from L. aurea (n=4), and each cryoprotectant (15% v/v DMSO with either 1 or 10% w/v sucrose) both 0- and 30-minutes post thaw. Estimated marginal means determined by generalised linear mixed modelling. Error bars = 95% confidence intervals. (a) Proportion motile (b) Proportion live.

### 3.3 Experiment 3: Cold storage Study

The predicted decline in motility was steady over the 14 day period (b=-0.193, t(1)=-7.5, p<0.001). Predicted motility began at approximately 80% and reduced to approximately 20% over a 14-day period. The GLMM model predicted motility declines to 50% by 7-8 days (Figure 4).

**Figure 4.**
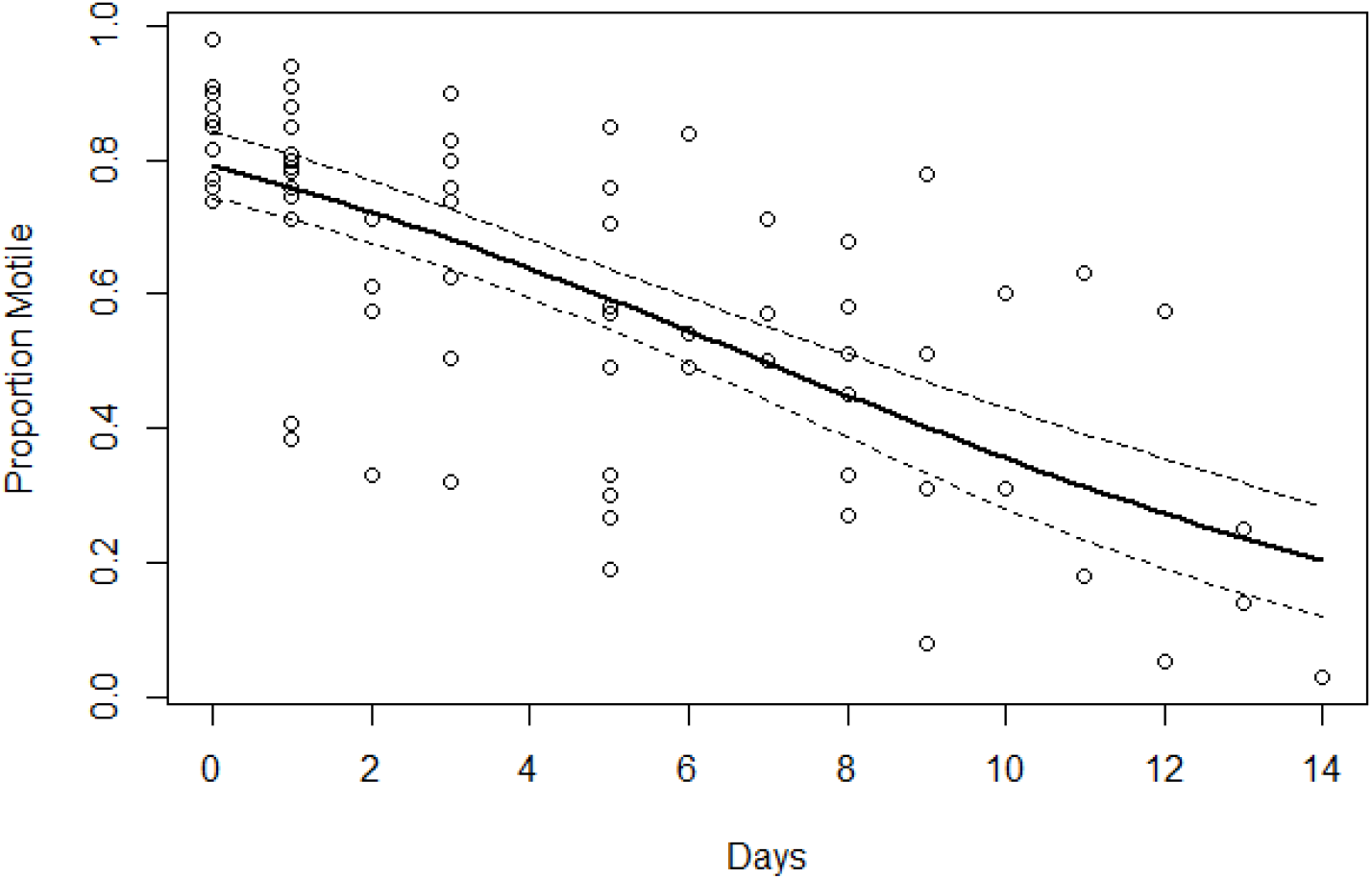
Reduction of motility of urinic sperm samples of L. aurea (n=7) stored at 5°C. Graph depicts raw data (○) with the predicted average decline in motility (black line) determined from a generalised linear model. Dotted lines = 95% confidence band.

## 4. Discussion

This study adds to a growing number of successful protocols used to induce non-lethal release of sperm from the testes of amphibians by injection of exogenous hormones, demonstrating this approach to obtaining male gametes for ART is a widely applicable procedure across many species. Nevertheless, there are fewer reports of the induction protocols being extended to the next phase of ART development, namely extended cold storage of urinic sperm without freezing [18, 27, 29, 34, 36, 46], and by cryopreservation [23, 24]. This study has demonstrated that all three procedures are possible in a threatened frog including sperm induction with exogenous hormones, retention of motility when unfrozen sperm were stored at 5°C for up to 14 days, and strong recovery of motility and vitality post-thaw after cryopreservation. This largely completes the suite of technologies required for collecting, manipulating and storing male gametes in threatened species.

We found large differences in the sperm inducing power of the two hormonal regimes tested. Our study found GnRH-a relatively ineffective in inducing concentrated sperm numbers when compared to hCG. The hCG response was essentially saturated at 20 IU/g body weight, and actually declined marginally at 40 IU/g, although that decline was not significant. These data support reports of induced spermiation in other pelodryadid species using hCG induction in a range ∼60-300 IU per whole animal injection [14, 17, 19], indicating the powerful response of many amphibian species to this synthetic mammalian gonadotropin. A recent study by Silla et al. (2019) on a pelodryadid species, *Litoria booroolongensis*, found a similar response to hCG at 20 and 40 IU/g body weight with sperm release also peaking after 2-4 hours [17]. Also noteworthy, the current study was performed at the shoulder of the natural breeding season, indicating the induction of spermiation in *L. aurea* does not need to occur at the peak of the breeding season in this Australian temperate species, a feature which may extend to a range of other Australian temperate frogs which are known to call over extended periods of the year [14, 47].

The response to GnRH-a was also similar in the present study compared to both Silla et al. (2019) and Pham and Brannelly (2021). As with hCG, lower doses of GnRH-a induced higher numbers of sperm release in *L. aurea* (this study) and in *L. booroolongensis* [17], although the differences weren’t significant in the current study. The endocrinology of declining sperm release with higher doses of GnRH-a and hCG is not resolved in either study but indicates the importance of testing a number of doses to determine optimal dose-response curves when optimising such protocols. Such effects may be due to complex events such as receptor saturation, and the induction of feed-back loops [14]. Taken together, these data reinforce the view that hCG is likely to be the optimal choice for induction of spermiation across a range of pelodryadid species, in comparison to GnRH-a. Nevertheless, even though GnRH-a is less potent in spermiation, both spermiation and breeding behaviours have also been induced in the Southern bell frog (*L. raniformis*) using GnRH-a [48], a closely related species to *L. aurea*.

Of the few studies that examine the effects of cold storage on anuran sperm motility, only few have utilised urinic sperm [18, 27, 29, 34, 46]. Testicular macerates are in totally different ionic and osmotic environment to urinic sperm. This is important because sperm stored in the testes are usually macerated in isotonic media, such as SAR (∼220 mOsm/kg) in which sperm remain immotile until activation by hypoosmotic shock [49, 50]. Cold storage of osmotically inactivated sperm is thought to reduce metabolic rates and energy utilisation by the sperm cell, allowing storage for longer periods of time [4]. This has been the paradigm in studies of various amphibian species that utilise either whole testes or testicular macerates [11, 30, 31, 33-35, 51-53].

In contrast, urinic sperm undergoes a hypoosmotic shock during its release in urine and is thus motile from the commencement of storage. In a range of Bufonid species (*Anaxyrus boreas boreas, A. fowleri* and *Peltophryne lemur*), urinic sperm has been observed to range from 20-80 mOsm/Kg and in *Lithobates sevosa*, 45 mOsm/Kg (Langhorne, 2016). While *a priori* that this may be expected to pose a problem for longevity of motility during storage, the present study found that urinic sperm of *L. aurea* can retain motility for up to 14 days at 5°C with the predicted drop to 50% motile sperm at 7 days. Similar studies utilising urinic sperm have found retention of ∼12-50% of motility in three species, after 2-6 days at 4-5°C [27, 29, 34]. While these results suggest some variation in preservation of motility for different species, taken together, the storage of both testicular and urinic sperm at low temperatures for later use in IVF or for cryopreservation is probably feasible in a wide array of species.

There are only four full reports of cryopreservation of urinic sperm in the literature [23-25, 27], an unpublished thesis [54] and a brief report [55]. Those species were from the families Ranidae and Bufonidae (not closely related to pelodryadid species) and produced large enough volumes of urinic sperm per animal to readily test and replicate a range of cryopreservation protocols, typically utilising either DMSO or N,N-dimethylformamide (DMFA) [24, 54].

In a previous study using testicular macerates, we noted post-thaw sperm vitality was higher than post-thaw motility, indicative of non-lethal cryopreservation related injury to sperm [39]. In the present study, this relationship remained true for the cryoprotectant previously used in this species (15% v/v DMSO with 10% w/v sucrose), but not true for the newly trialled cryodiluent, 15% v/v DMSO with only 1% w/v sucrose. This result indicates that the lower concentration of non-penetrating cryoprotectant (sucrose) in the cryodiluent minimises non-lethal damage and also improves on the previous method by a significant margin in terms of recovery of viable and motile sperm after cryopreservation.

This study further considered the practicality of using cryopreserved sperm for IVF by assessing post-thaw motility at both 0- and 30-minutes post-thaw. While there was a slight decrease in both motility and vitality 30 minutes post-thaw in both treatments, this decrease was not significant. The results suggest a user window following thawing of sperm would be at least half an hour, which should allow time for post-thaw assessments, adjustment of sperm concentration, wash steps and addition of sperm to ova for fertilisation purposes.

Additionally, the ability to easily induce a spermiation response in males followed by storage at 0-5°C or cryopreservation provides protocols that indicate that non-lethal collection and storage of sperm could be successfully combined with fertilisation of ova when these are available.

## 5. Conclusions

Our study shows that sperm release can be induced in high concentrations with 20 IU/g bodyweight of human chorionic gonadotrophin (hCG). For the purpose of storing sperm we demonstrated that high levels of live, motile sperm could be recovered post-cryopreservation by treating the sperm with 15% v/v DMSO and 1% w/v sucrose pre-freeze and that urinic sperm could be stored at 5°C retaining motility over a 14-day period. Our findings demonstrate that it is possible to obtain and store large quantities of quality sperm from a threatened amphibian via non-lethal means, representing an important step forward for the use of ARTs in conservation programs for rare and threatened species.

## 8. Author Contributions

RU, NEC, SC, MM and JC conceived the study and designed the experiments. RU provided husbandry for all animals involved in the study. RU, NEC and AB performed hormone induction experiments; RU and DB conducted cryopreservation experiments; RU and JC conducted short-term storage experiments. RU and KC statistically analysed the data. RU wrote the paper with input from all other authors.

## 9. Acknowledgements

The authors wish to thank Justine O’Brien and Rebecca Hobbs for provision of laboratory resources and Alex Callen and Jasmine Callen for assistance maintaining *L. aurea*. We would also like to acknowledge and respect the Pambalong people of the Awabakal Nation, the traditional custodians of the land on which we work, and pay respect to the Elders past, present and future.

## References

[1] Gillespie GR, Roberts JD, Hunter D, Hoskin CJ, Alford RA, Heard GW, et al. Status and priority conservation actions for Australian frog species. Biol Conserv. 2020;247:108543.

[2] Stuart SN, Chanson JS, Cox NA, Young BE, Rodrigues ASL, Fischman DL, et al. Status and trends of amphibian declines and extinctions worldwide. Science. 2004;306:1783–6.

[3] Beranek CT, Hamer AJ, Mahony SV, Stauber A, Ryan SA, Gould J, et al. Severe wildfires promoted by climate change negatively impact forest amphibian metacommunities. Diversity and Distributions. 2023;29:785–800.

[4] Browne RK, Silla AJ, Upton R, Della-Togna G, Marcec-Greaves R, Shishova NV, et al. Sperm collection and storage for the sustainable management of amphibian biodiversity. Theriogenology. 2019;133:187–200.

[5] Clulow J, Upton R, Trudeau VL, Clulow S. Amphibian Assisted Reproductive Technologies: Moving from Technology to Application. In: Comizzoli P, Brown JL, Holt WV, editors. Reproductive Sciences in Animal Conservation. Cham: Springer International Publishing; 2019. p. 413–63.

[6] Della Togna G, Howell LG, Clulow J, Langhorne CJ, Marcec-Greaves R, Calatayud NE. Evaluating amphibian biobanking and reproduction for captive breeding programs according to the Amphibian Conservation Action Plan objectives. Theriogenology. 2020;150:412–31.

[7] Howell LG, Frankham R, Rodger JC, Witt RR, Clulow S, Upton RMO, et al. Integrating biobanking minimises inbreeding and produces significant cost benefits for a threatened frog captive breeding programme. Conservation Letters. 2020;e12776.

[8] Howell LG, Mawson PR, Frankham R, Rodger JC, Upton RM, Witt RR, et al. Integrating biobanking could produce significant cost benefits and minimise inbreeding for Australian amphibian captive breeding programs. Reprod Fertil Dev. 2021;33:573–87.

[9] Clulow S, Clulow J, Marcec-Greaves R, Della Togna G, Calatayud NE, Yuan Y. Common goals, different stages: the state of the ARTs for reptile and amphibian conservation. Reprod Fertil Dev. 2022;34:i–ix.

[10] Browne RK, Clulow J, Mahony MJ, Clark AK. Successful recovery of motility and fertility of cryopreserved cane toad (Bufo marinus) sperm. Cryobiology. 1998;37:339–45.

[11] Browne RK, Clulow J, Mahony MJ. The short-term storage and cryopreservation of spermatozoa from hylid and myobatrachid frogs. Cryoletters. 2002;23:129–36.

[12] Mugnano JA, Costanzo JP, Beesley SG, Lee RE. Evaluation of glycerol and dimethyl sulfoxide for the cryopreservation of spermatozoa from the wood frog (Rana sylvatica). Cryoletters. 1998;19:249–54.

[13] Upton R, Clulow S, Mahony MJ, Clulow J. Generation of a sexually mature individual of the Eastern dwarf tree frog, Litoria fallax, from cryopreserved testicular macerates: proof of capacity of cryopreserved sperm derived offspring to complete development. Conserv Physiol. 2018;6:coy043–coy.

[14] Clulow J, Pomering M, Herbert D, Upton R, Calatayud N, Clulow S, et al. Differential success in obtaining gametes between male and female Australian temperate frogs by hormonal induction: A review. Gen Comp Endocrinol. 2018.

[15] Kouba AJ, delBarco-Trillo J, Vance CK, Milam C, Carr M. A comparison of human chorionic gonadotropin and luteinizing hormone releasing hormone on the induction of spermiation and amplexus in the American toad (Anaxyrus americanus). Reproductive Biology and Endocrinology : RB&E. 2012;10:59-.

[16] Silla AJ, Roberts JD. Investigating patterns in the spermiation response of eight Australian frogs administered human chorionic gonadotropin (hCG) and luteinizing hormone-releasing hormone (LHRHa). Gen Comp Endocrinol. 2012;179:128–36.

[17] Silla AJ, McFadden MS, Byrne PG. Hormone-induced sperm-release in the critically endangered Booroolong frog (Litoria booroolongensis): effects of gonadotropin-releasing hormone and human chorionic gonadotropin. Conserv Physiol. 2019;7.

[18] Arregui L, Diaz-Diaz S, Alonso-López E, Kouba AJ. Hormonal induction of spermiation in a Eurasian bufonid (Epidalea calamita). Reprod Biol Endocrinol. 2019;17:92.

[19] Pham TH, Brannelly LA. Sperm parameters following hormonal induction of spermiation in an endangered frog [the alpine tree frog] (Litoria verreauxii alpina). Reprod Fertil Dev. 2022;34:867–74.

[20] Della Togna G, Trudeau VL, Gratwicke B, Evans M, Augustine L, Chia H, et al. Effects of hormonal stimulation on the concentration and quality of excreted spermatozoa in the critically endangered Panamanian golden frog (Atelopus zeteki). Theriogenology. 2017;91:27–35.

[21] Trudeau VL, Somoza GM, Natale GS, Pauli B, Wignall J, Jackman P, et al. Hormonal induction of spawning in 4 species of frogs by coinjection with a gonadotropin-releasing hormone agonist and a dopamine antagonist. Reprod Biol Endocrinol. 2010;8:1–9.

[22] Vu M, Weiler B, Trudeau VL. Time- and dose-related effects of a gonadotropin-releasing hormone agonist and dopamine antagonist on reproduction in the Northern leopard frog (Lithobates pipiens). Gen Comp Endocrinol. 2017;254:86–96.

[23] Shishova NR, Uteshev VK, Kaurova SA, Browne RK, Gakhova EN. Cryopreservation of hormonally induced sperm for the conservation of threatened amphibians with Rana temporaria as a model research species. Theriogenology. 2011;75:220–32.

[24] Uteshev VK, Shishova N, Kaurova S, Manokhin A, Gakhova E. Collection and cryopreservation of hormonally induced sperm of pool frog (Pelophylax lessonae). Russ J Herpetol. 2013;20:105–9.

[25] Hinkson KM, Baecher JA, Poo S. Cryopreservation and hormonal induction of spermic urine in a novel species: The smooth-sided toad (Rhaebo guttatus). Cryobiology. 2019;89:109–11.

[26] Burger IJ, Lampert SS, Kouba CK, Morin DJ, Kouba AJ. Development of an amphibian sperm biobanking protocol for genetic management and population sustainability. Conserv Physiol. 2022;10.

[27] Arregui L, Bóveda P, Gosálvez J, Kouba AJ. Effect of seasonality on hormonally induced sperm in Epidalea calamita (Amphibia, Anura, Bufonidae) and its refrigerated and cryopreservated storage. Aquaculture. 2020;529:735677.

[28] Guy EL, Gillis AB, Kouba AJ, Barber D, Poole V, Marcec-Greaves RM, et al. Sperm collection and cryopreservation for threatened newt species. Cryobiology. 2020;94:80–8.

[29] Germano JM, Arregui L, Kouba AJ. Effects of aeration and antibiotics on short-term storage of Fowler’s toad (Bufo fowleri) sperm. Aquaculture. 2013;396–399:20–4.

[30] Browne R, Clulow J, Mahony M. Short-term storage of cane toad (Bufo marinus) gametes. Reproduction. 2001;121:167–73.

[31] Browne RK, Davis J, Clulow J, Pomering M. Storage of cane toad Bufo marinus sperm for 6 days at 0°C with subsequent cryopreservation. Reprod Fertil Dev. 2002;14:267–73.

[32] Figiel CR. Cold storage of sperm from the axolotl, Ambystoma mexicanum. Herpetol Conserv Biol. 2020;15:367–71.

[33] Silla AJ, Keogh LM, Byrne PG. Antibiotics and oxygen availability affect the short-term storage of spermatozoa from the critically endangered booroolong frog, Litoria booroolongensis. Reprod Fertil Dev. 2015.

[34] Keogh LM, Byrne PG, Silla AJ. The effect of gentamicin on sperm motility and bacterial abundance during chilled sperm storage in the Booroolong frog. Gen Comp Endocrinol. 2017;243:51–9.

[35] Shishova NV, Uteshev VK, Sirota NP, Kuznetsova EA, Kaurova SA, Browne RK, et al. The quality and fertility of sperm collected from European common frog (Rana temporaria) carcasses refrigerated for up to 7 days. Zoo Biol. 2013;32:400–6.

[36] Uteshev V, Kaurova S, Shishova N, Stolyarov S, Browne R, Gakhova E. In vitro Fertilization with hormonally induced sperm and eggs from sharp-ribbed newts, Pleurodeles waltl. Russ J Herpetol. 2015;22.

[37] Goncharov BF, Shubravy OI, Serbinova IA, Uteshev VK. The USSR programme for breeding amphibians, including rare and endangered species. Int Zoo Yearb. 1989;28:10–21.

[38] Mahony MJ, Hamer AJ, Pickett EJ, McKenzie DJ, Stockwell MP, Garnham JI, et al. Identifying conservation and research priorities in the face of uncertainty: a review of the threatened bell frog complex in eastern Australia. Herpetol Conserv Biol. 2013;8:519–38.

[39] Upton R, Clulow S, Calatayud NE, Colyvas K, Seeto RG, Wong LA, et al. Generation of reproductively mature offspring from the endangered green and golden bell frog Litoria aurea using cryopreserved spermatozoa. Reprod Fertil Dev. 2021;33:562–72.

[40] Upton R, Clulow S, Colyvas K, Mahony M, Clulow J. Paradigm shift in frog sperm cryopreservation: reduced role for non-penetrating cryoprotectants. Reproduction. 2023;165:583–92.

[41] Brooks ME, Kristensen K, van Benthem KJ, Magnusson A, Berg CW, Nielsen A, et al. glmmTMB balances speed and flexibility among packages for zero-inflated generalized linear mixed modeling. The R journal. 2017;9:378–400.

[42] Bates D, Mächler M, Bolker B, Walker S. Fitting Linear Mixed-Effects Models Using lme4. J Stat Softw. 2015;67:1–48.

[43] Lenth R, Singmann H, Love J, Buerkner P, Herve M. Emmeans: Estimated marginal means, aka least-squares means. R package version 122: https://CRAN.R-project.org/package=emmeans; 2018.

[44] Auguie B, Antonov A. gridExtra: miscellaneous functions for “grid” graphics. R package version. 2017;2:602.

[45] Wickham H. ggplot2: elegant graphics for data analysis: springer; 2016.

[46] Langhorne CJ, Calatayud NE, Kouba CK, Willard ST, Smith T, Ryan PL, et al. Efficacy of hormone stimulation on sperm production in an alpine amphibian (Anaxyrus boreas boreas) and the impact of short-term storage on sperm quality. Zoology (Jena). 2021;146:125912.

[47] Lemckert F, Mahony M. Core calling periods of the frogs of temperate New South Wales, Australia. Herpetol Conserv Biol. 2008;3:71–6.

[48] Mann RM, Hyne RV, Choung CB. Hormonal induction of spermiation, courting behavior and spawning in the southern bell frog, Litoria raniformis. Zoo Biol. 2010;29:774–82.

[49] Inoda T, Morisawa M. Effect of osmolality on the initiation of sperm motility in Xenopus laevis. Comparative Biochemistry and Physiology Part A: Molecular & Integrative Physiology. 1987;88:539–42.

[50] Edwards DL, Mahony MJ, Clulow J. Effect of sperm concentration, medium osmolality, and oocyte storage on artificial fertilisation success in a myobatrachid frog (Limnodynastes tasmaniensis) Reprod Fertil Dev. 2004;16:347–54.

[51] Rostand J. Glyérine et reistance du sperme aux basses températures. Comptes Rendus de l’Académie des Sciences (Paris). 1946;222:1524–5.

[52] Rostand J. Sur le refroidissement des cellules spermatiques en presénce de glycérine. Comptes Rendus Hebdomadaires Des Seances de L’Academie des Sciences. 1952;234:2310–2.

[53] Silla AJ. Artificial fertilisation in a terrestrial toadlet (Pseudophryne guentheri): effect of medium osmolality, sperm concentration and gamete storage. Reprod Fertil Dev. 2013;25:1134–41.

[54] Langhorne CJ. Developing Assisted Reproductive Technologies for Endangered North American Amphibians: Mississippi State University; 2016.

[55] Lampert SS, Burger IJ, Julien AR, Gillis AB, Kouba AJ, Barber D, et al. Sperm Cryopreservation as a Tool for Amphibian Conservation: Production of F2 Generation Offspring from Cryo-Produced F1 Progeny. Animals. 2023;13:53.

