## Supplementary Materials for "Refrigerated storage and cryopreservation of hormone induced sperm in a threatened frog"

*Supplementary Table 1. Ratios and 95% confidence intervals for pairwise comparisons of 4 hormone treatments (20 and 40 IU/g bodyweight hCG and 0.25 and 0.5 µg/g bodyweight GnRH-a) to compare induction of sperm concentration (cells/mL) in* L. aurea *(n=5 per treatment), averaged across all collection times.*

| Pairwise Comparison (treatment) | | Ratio | LCL^3^ | UCL^4^ |
| --- | --- | --- | --- | --- |
| 20 IU/g hCG^1^ | 40 IU/g hCG | 1.3 | 0.2 | 7.9 |
| 20 IU/g hCG | 0.25 µg/g GnRH-a | 14.3 | 2.3 | 89.3 |
| 20 IU/g hCG | 0.5 µg/g GnRH-a | 36.7 | 5.8 | 232.3 |
| 40 IU/g hCG | 0.25 µg/g GnRH-a | 11.2 | 1.8 | 70.3 |
| 40 IU/g hCG | 0.5 µg/g GnRH-a | 28.6 | 4.5 | 182.4 |
| 0.25 µg/g GnRH-a^2^ | 0.5 µg/g GnRH-a | 2.6 | 0.4 | 16.0 |

^1^Human chorionic gonadotrophin

^2^Gonadotrophin-releasing hormone

^3^Lower confidence limit

^4^Upper confidence limit

*Supplementary Table 2. Ratios and 95% confidence intervals for pairwise comparisons of 4 hormone treatments (20 and 40 IU/g bodyweight hCG and 0.25 and 0.5 µg/g bodyweight GnRH-a) to compare induction of total sperm numbers (cells/sample) in* L. aurea *(n=5 per treatment), averaged across all collection times.*

| Pairwise Comparison (treatment) | | Ratio | LCL^3^ | UCL^4^ |
| --- | --- | --- | --- | --- |
| 20 IU/g hCG^1^ | 40 IU/g hCG | 1.1 | 0.2 | 6.9 |
| 20 IU/g hCG | 0.25 µg/g GnRH-a | 15.6 | 2.6 | 94.9 |
| 20 IU/g hCG | 0.5 µg/g GnRH-a | 20.3 | 3.3 | 126.0 |
| 40 IU/g hCG | 0.25 µg/g GnRH-a | 13.8 | 2.2 | 85.5 |
| 40 IU/g hCG | 0.5 µg/g GnRH-a | 17.9 | 2.9 | 112.2 |
| 0.25 µg/g GnRH-a^2^ | 0.5 µg/g GnRH-a | 1.3 | 0.2 | 8.0 |

^1^Human chorionic gonadotrophin

^2^Gonadotrophin-releasing hormone

^3^Lower confidence limit

^4^Upper confidence limit

*Supplementary Table 3. Ratios and 95% confidence intervals for pairwise comparisons of 4 hormone treatments (20 and 40 IU/g bodyweight hCG and 0.25 and 0.5 µg/g bodyweight GnRH-a) to compare collection hour in* L. aurea *(n=5 per treatment), averaged across all treatments.*

| Pairwise Comparison (Collection hour) | | Sperm Concentration | | | Total Sperm | | |
| --- | --- | --- | --- | --- | --- | --- | --- |
|  |  | Ratio | LCL^1^ | UCL^2^ | Ratio | LCL | UCL |
| 1 | 2 | 0.9 | 0.5 | 1.7 | 0.6 | 0.3 | 1.2 |
| 1 | 3 | 0.8 | 0.4 | 1.6 | 1.0 | 0.5 | 2.2 |
| 1 | 4 | 1.5 | 0.7 | 3.4 | 2.5 | 1.1 | 5.8 |
| 1 | 5 | 1.9 | 0.9 | 4.0 | 4.8 | 2.0 | 11.3 |
| 1 | 6 | 3.2 | 1.5 | 7.0 | 7.4 | 3.3 | 16.9 |
| 1 | 24 | 4.5 | 1.9 | 10.7 | 13.1 | 4.9 | 34.9 |
| 1 | 48 | 13.8 | 4.6 | 41.6 | 44.4 | 12.5 | 157.9 |
| 2 | 3 | 0.9 | 0.5 | 1.7 | 1.7 | 0.9 | 3.5 |
| 2 | 4 | 1.7 | 0.8 | 3.6 | 4.2 | 2.0 | 8.9 |
| 2 | 5 | 2.1 | 1.0 | 4.2 | 7.9 | 3.6 | 17.5 |
| 2 | 6 | 3.5 | 1.7 | 7.3 | 12.3 | 5.8 | 26.2 |
| 2 | 24 | 5.0 | 2.2 | 11.3 | 21.8 | 8.7 | 54.5 |
| 2 | 48 | 15.3 | 5.3 | 44.1 | 73.8 | 21.3 | 256.3 |
| 3 | 4 | 1.9 | 1.0 | 3.9 | 2.4 | 1.1 | 5.2 |
| 3 | 5 | 2.4 | 1.2 | 4.5 | 4.6 | 2.2 | 9.6 |
| 3 | 6 | 4.0 | 2.1 | 7.7 | 7.1 | 3.5 | 14.6 |
| 3 | 24 | 5.7 | 2.6 | 12.4 | 12.5 | 5.1 | 30.7 |
| 3 | 48 | 17.3 | 6.1 | 48.5 | 42.5 | 12.4 | 145.4 |
| 4 | 5 | 1.2 | 0.6 | 2.6 | 1.9 | 0.8 | 4.3 |
| 4 | 6 | 2.1 | 1.0 | 4.4 | 2.9 | 1.3 | 6.4 |
| 4 | 24 | 2.9 | 1.3 | 6.8 | 5.2 | 2.0 | 13.2 |
| 4 | 48 | 8.9 | 3.1 | 25.5 | 17.5 | 5.1 | 60.8 |
| 5 | 6 | 1.7 | 0.8 | 3.4 | 1.6 | 0.7 | 3.3 |
| 5 | 24 | 2.4 | 1.1 | 5.4 | 2.7 | 1.1 | 6.8 |
| 5 | 48 | 7.3 | 2.5 | 21.5 | 9.3 | 2.6 | 33.2 |
| 6 | 24 | 1.4 | 0.6 | 3.2 | 1.8 | 0.8 | 4.1 |
| 6 | 48 | 4.3 | 1.5 | 12.3 | 6.0 | 1.8 | 19.8 |
| 24 | 48 | 3.0 | 1.0 | 9.3 | 3.4 | 0.9 | 12.2 |

^1^Lower confidence limit

^2^Upper confidence limit

*Supplementary Table 4. Odds ratios and 95% confidence intervals for pairwise comparisons of 4 hormone treatments (20 and 40 IU/g bodyweight hCG and 0.25 and 0.5 µg/g bodyweight GnRH-a) to compare motility of urinic sperm in* L. aurea *(n=5 per treatment), averaged across all collection time points.*

| Pairwise Comparison | | Odds Ratio | LCL^3^ | UCL^4^ |
| --- | --- | --- | --- | --- |
| 20 IU/g hCG^1^ | 40 IU/g hCG | 5.0 | 1.8 | 13.4 |
| 20 IU/g hCG | 0.5 µg/g GnRH | 11.7 | 4.1 | 33.4 |
| 20 IU/g hCG | 0.25 µg/g GnRH | 3.5 | 1.2 | 9.8 |
| 40 IU/g hCG | 0.5 µg/g GnRH | 2.3 | 0.8 | 6.8 |
| 0.25 µg/g GnRH^2^ | 40 IU/g hCG | 1.4 | 0.5 | 4.0 |
| 0.25 µg/g GnRH | 0.5 µg/g GnRH | 3.4 | 1.1 | 10.0 |

^1^Human chorionic gonadotrophin

^2^Gonadotrophin-releasing hormone

^3^Lower confidence limit

^4^Upper confidence limit
